# Circadian rhythm disruption alters rhythmic properties of clock and memory gene expression in the hippocampi of rats

**DOI:** 10.64898/2026.07.15.738771

**Authors:** Scott H. Deibel, Andrew B. Lehr, Stavroula Michalopoulou, Incé Husain, Eugenio F. Fornasiero, Cameron Bye, Nancy S. Hong, Olga Kovalchuk, Robert J. McDonald

**Affiliations:** Canadian Centre for Behavioural Neuroscience, Department of Neuroscience, University of Lethbridge, Lethbridge, Alberta, Canada; Department of Neuro- and Sensory Physiology, University Medical Center Göttingen, Göttingen, Germany; Circulant Labs, Bensheim, Germany; University of New Brunswick, Department of Psychology, Fredericton, New Brunswick, Canada; Department of Life Sciences, University of Trieste, Trieste, Italy

**Author notes:** Both authors contributed equally to this work (Scott H. Deibel, Andrew B. Lehr). Corresponding authors: Scott H. Deibel Department of Neuroscience University of Lethbridge Lethbridge, Alberta, Canada T1K 3M4, Robert J. McDonald Department of Neuroscience University of Lethbridge Lethbridge, Alberta, Canada T1K 3M4.

**Keywords:** Circadian rhythm disruption, forced desynchrony, health, memory, gene expression, rats

## Abstract

Our modern environment – with its artificial lighting, irregular work hours, and frequent travel — often disrupts our circadian rhythms, which can lead to health problems, particularly in learning and memory. This is especially concerning given the aging population and the rising prevalence of dementia. Yet, the biological mechanisms linking circadian disruption to cognitive impairment remain poorly understood. At the molecular level, genetic techniques have been used to attenuate or abolish expression of key genes involved in circadian rhythms and these manipulations have detrimental effects on memory function. However, whether environmentally induced circadian disruption, impairs memory via changes in overall gene expression levels in the hippocampus or rather via changes in the coordinated rhythmic patterns of circadian expression across groups of genes is less known. Here, we examined how environmental circadian disruption affects the expression of genes involved in the circadian clock and memory in the hippocampus of rats using a forced desynchrony model. Circadian disruption changed the rhythmic properties of gene expression in most genes assessed but had no measurable effect on average expression levels across the day. These findings suggest that the inability to maintain circadian synchrony rather than overall expression may underlie the cognitive deficits observed in circadian-related disorders.

## Introduction

Circadian rhythms are 24 hour cycles in biological processes or behavior that allow organisms to synchronize with their environment ^1–5^. These are generated at the cellular level by translation/transcription feedback loops of certain genes – *clock* genes – that take approximately 24 hours ^6–9^. Circadian rhythms are involved in virtually all aspects of health and disease such as cognition, aging, cardiovascular disease, metabolic diseases, cancer, mental illness, and stroke ^2,4,5,10^. Problems arise when the many processes in the brain and body that are under full or partial circadian control become desynchronized from each other or the environment ^2–5,10^. Human circadian rhythm disruption in its most common forms, night work, shift work, jetlag, or social-jetlag ^2,3,10^, involves a decoupling of one’s circadian rhythms from the environmental cues (zeitgebers) that set them.

There is a long history investigating the association between memory and circadian rhythms ^11–13^. While the relationship has been established the mechanisms at the cellular, synaptic, and network level are largely unknown^11^. Genetic manipulations in mice have unearthed much of what we know about this relationship^11,13,14^. For example, signal transduction pathways in the hippocampus important for memory consolidation are affected by *clock* gene knockouts and manipulating *clock* genes causes memory impairments^11,13,14^. Focusing on a gene or gene in a specific brain region has the advantage of isolating its specific effect on circadian rhythm function. However, studying circadian rhythm disruption similar to that which is thought to occur in humans depends on environmental manipulations of zeitgebers instead, in well-established translational animal models.

In the laboratory, forced desynchrony ^15–19^ is an established model that creates circadian rhythm disruption by employing a day length to which the organisms master clock in the brain – suprachiasmatic nucleus (SCN) – cannot synchronize. Importantly, activity rhythms are still oscillating in a circadian manner, but the animals’ internal timekeepers are decoupled from the primary zeitgebers in the environment (e.g. light) that they rely on to set the clocks^20^. This is akin to most types of human circadian rhythm disruption which still involves oscillating circadian rhythms, they are just not synchronized with the environment ^2,3^, and memory is impaired^15–19^. In particular, a number of studies suggest that attempting to learn a task during or after forced-desynchrony is made difficult by the detrimental effect of environmental circadian disruption on hippocampal dependent memory consolidation^20–23^.

However, the mechanisms by which environmental circadian rhythm disruption by forced desynchrony impacts gene expression are not well understood. To address this gap, we mapped the 24-hour expression profiles of a comprehensive set of hippocampal clock and memory genes in rats following a well-established acute forced desynchrony paradigm that disrupts learning and memory^24^. We demonstrate that while average gene expression remained unaffected, environmental circadian rhythm disruption severely impacted the temporal organization of hippocampal gene expression. Half of the genes that were circadian in control animals completely lost rhythmicity, the amplitude of oscillations was reduced, and the coordinated phase relationship between genes was destroyed. Ultimately, these findings reveal that cognitive deficits due to environmental circadian disruption may not stem from a lack of expression of key genes, as in gene knockout studies, but instead from a breakdown in molecular timing.

## Results

As depicted in Figure 1, after six days of exposure to either a 12L:12D light dark cycle (control: n = 24), or a 9L:12D (21-hr day) light dark cycle (shifted: n = 24) the whole hippocampus was extracted from eight animals every four hours (control: n = 4; shifted: n = 4) and the entire genome was assessed with next generation RNA sequencing. We analyzed a set of 11 clock and 18 memory-related genes for which recent studies have shown effects on memory when they are genetically or environmentally manipulated.^11,13,14,25–48^ (refer to Table 1 for genes selected).

**Figure 1.**
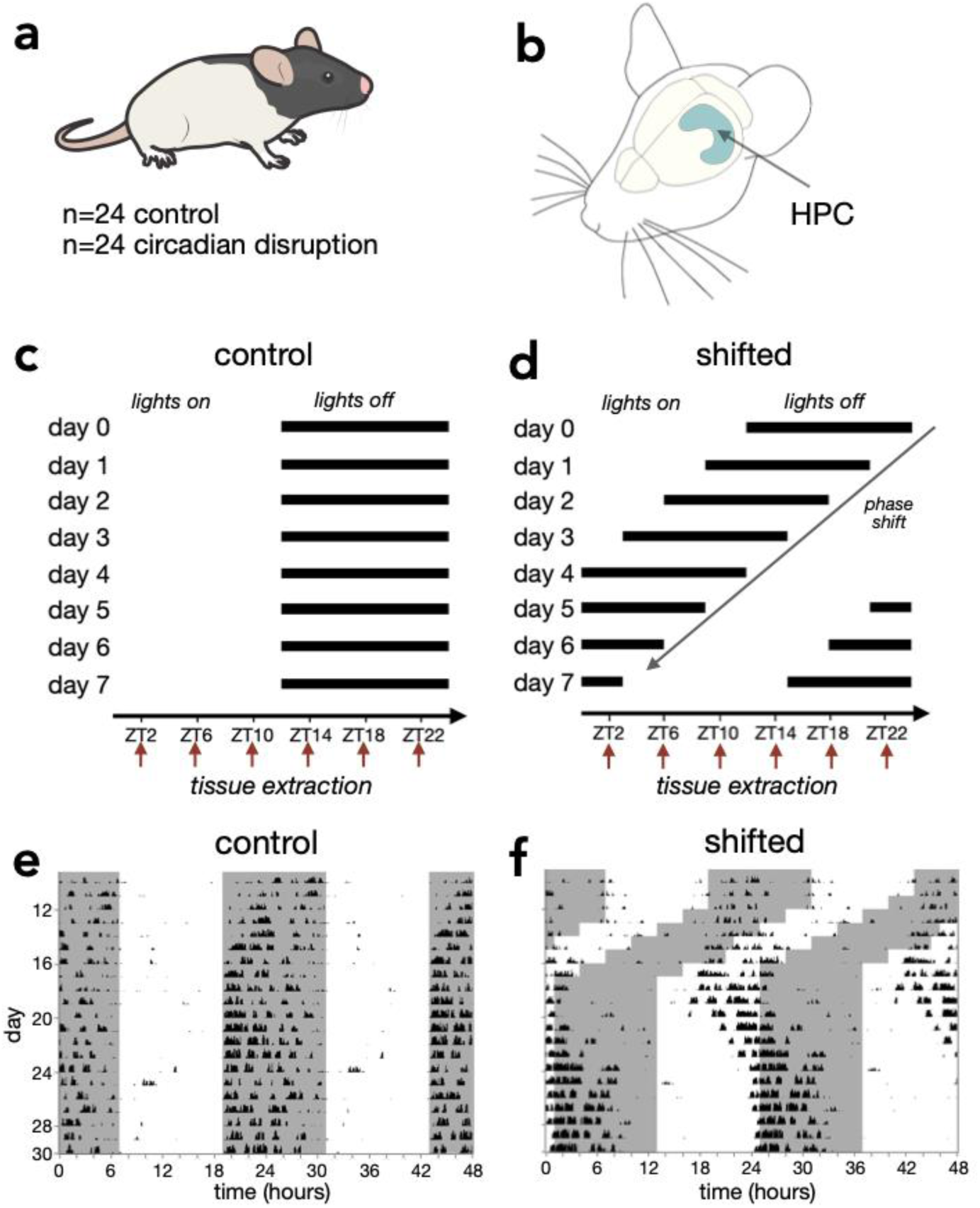
Measuring hippocampal gene expression after environmental circadian rhythm disruption using forced desynchrony. (a-d) The whole hippocampus was extracted every four hours from Long-Evans rats on day seven after six days of exposure to normal 12L:12D (n = 24; n = 4 per timepoint) and unsynchronizable (unentrainable) 9L:12D light-dark cycles (n = 24; n = 4 per timepoint). (e) Example actogram showing entrainment of wheel running activity to the normal 12L:12:D light-dark cycle. Lights on are indicated by white background, lights off by grey background. Data from Deibel and colleagues 2022, additional actograms in supplement. (f) Example actogram in the forced desynchrony paradigm showing stereotypical decoupling of activity from light-dark cycle. Note graphics in (a) from^49^ and (b) redrawn from^50^.

**Table 1.**
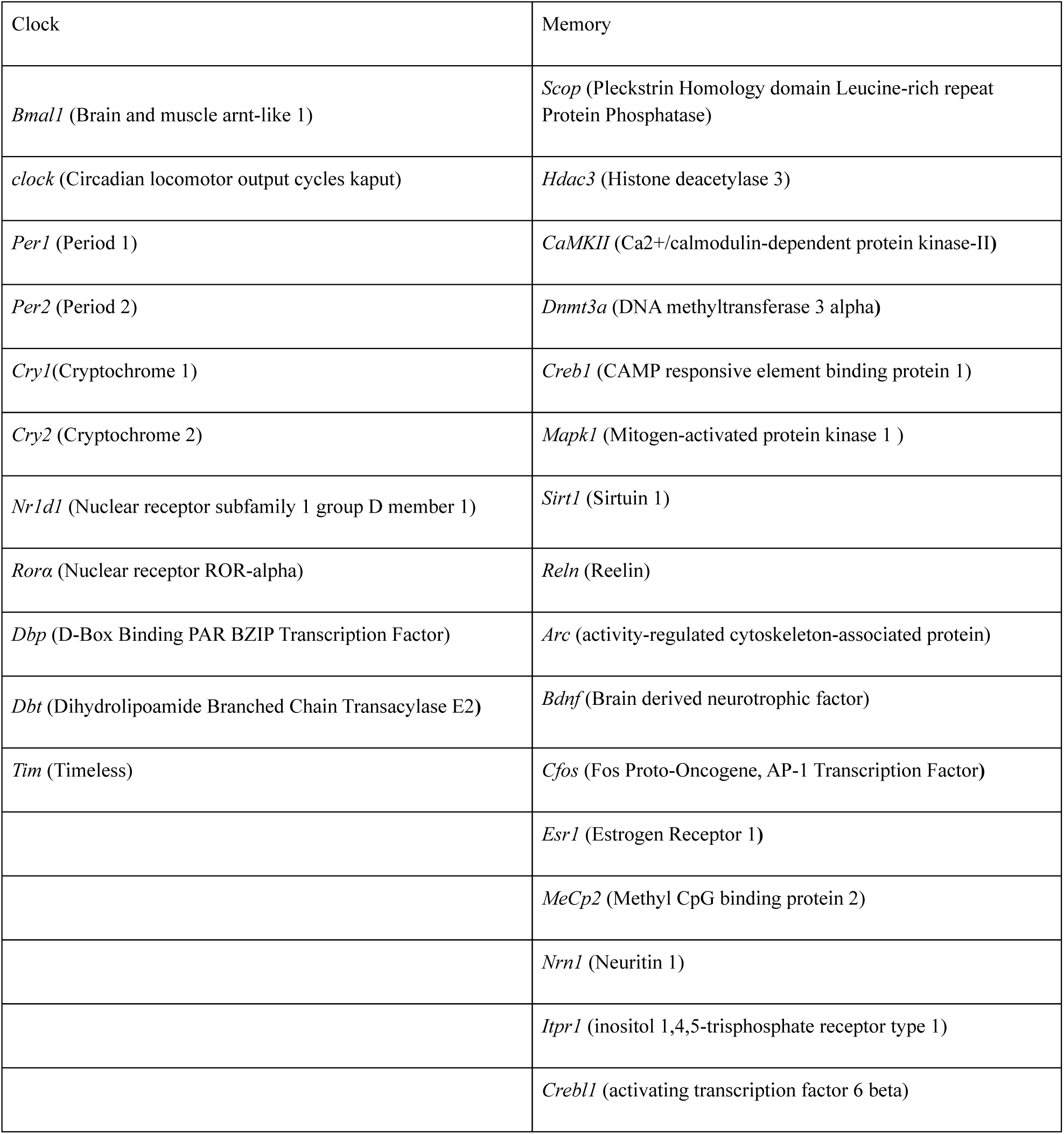
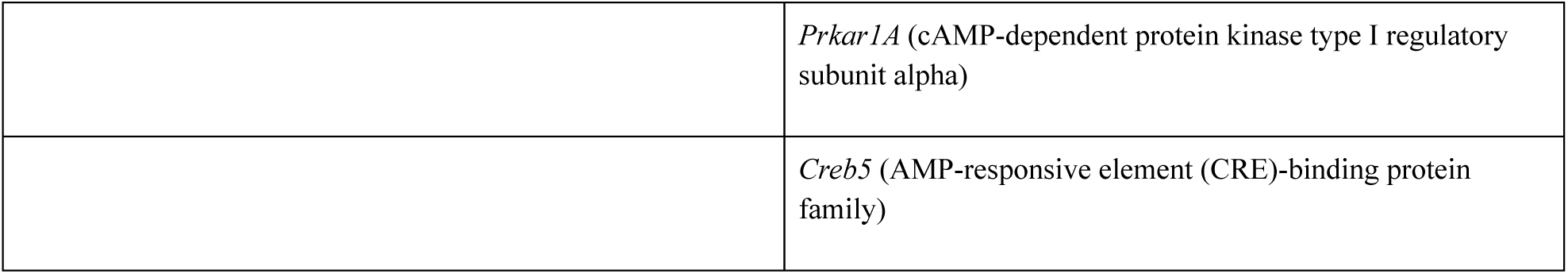
Genes involved in circadian rhythms and memory selected for analyses from a genome wide RNA sequencing dataset of whole rat hippocampus.

### Time-averaged gene expression is unchanged by circadian rhythm disruption

First, we were interested in whether circadian rhythm disruption changed the amount of gene expression averaged across all timepoints in each group (see Table 1 for a list of the genes analyzed). As shown in Figure 2a and supplementary Figure 1, we were not able to detect any up or down regulation for any of the genes tested. Despite amounts not changing, if genes display circadian rhythms in expression, it is possible that circadian rhythm disruption affects these oscillations. The next three analyses will study aspects of circadian rhythms in gene expression, first of which is whether the genes are expressed in a circadian manner under normal and disrupted lighting conditions.

**Figure 2.**
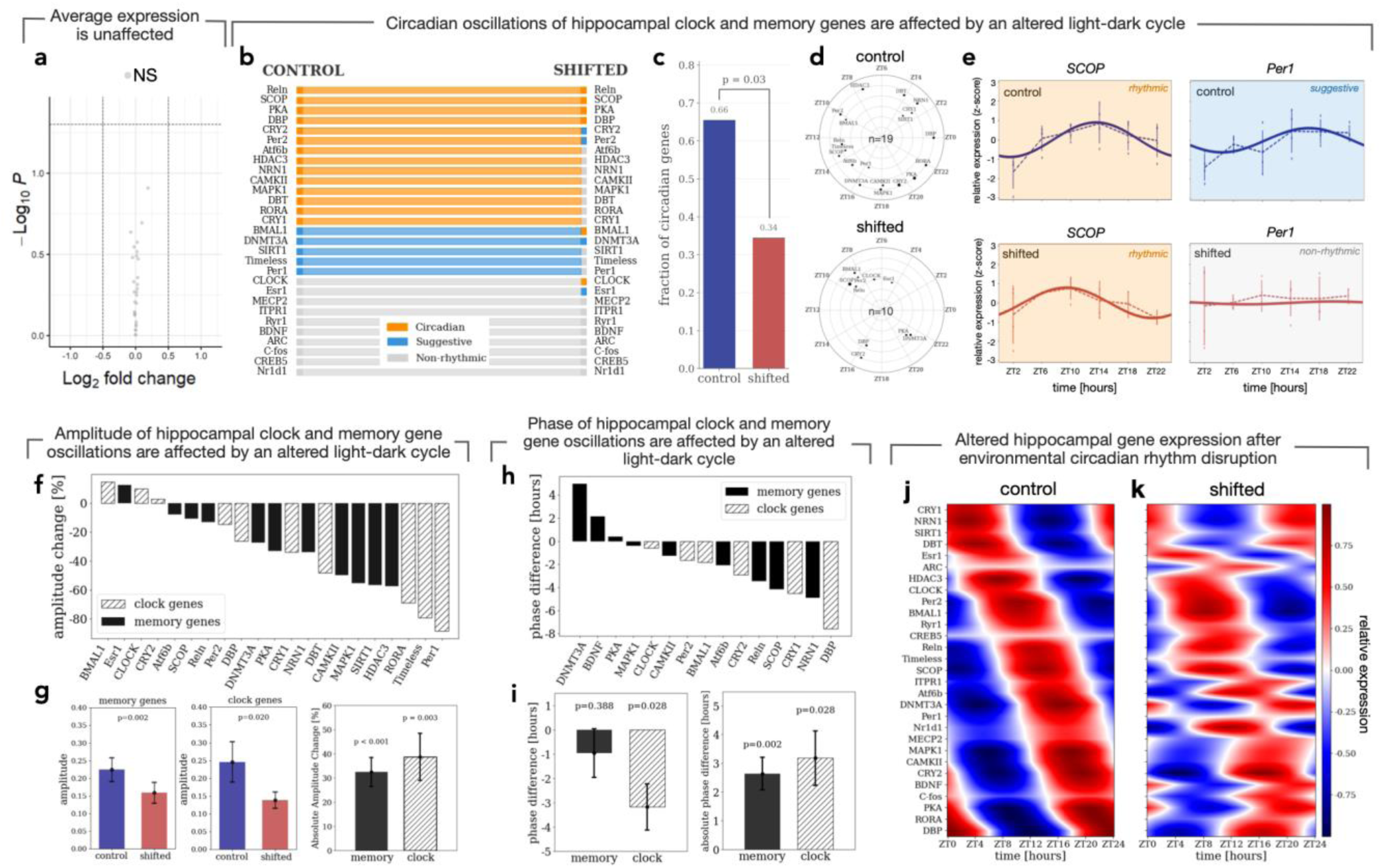
Circadian properties but not average expression are disrupted after environmental circadian disruption by forced desynchrony. (**a**) Gene expression averaged across timepoints was unaffected by circadian rhythm disruption, shown as volcano plot of statistical significance (-log10 *P*) versus change in expression (log2 fold change). (**b**) Alluvial plot identifying circadian, suggestive, or non-rhythmic genes as well as changes in circadian expression in control and phase shifted conditions. (**c**) Fraction of circadian genes (at least suggestive) in control and phase shifted conditions. (d) Polar plot showing the time of peak expression (angle) and amplitude of peak expression (radius). Relative size of point for each gene indicates significance level of circadian fit. Only suggestive and circadian genes are shown (control n=19 genes; shifted n=10 genes). (e) Examples of two genes in controls and phase shifted animals. *SCOP* was rhythmic in both controls and shifted animals, but with altered phase and amplitude of the rhythm. *Per1* was suggestive of circadian rhythms in controls but was no longer rhythmic after forced desynchrony. (f) Amplitude change (%) by gene. (g) Amplitude of expression is shown for control and phase shifted animals for memory (left) and clock (middle) genes and the absolute amplitude change is shown (right panel). (h) Difference in the time of peak expression (acrophase) between control and phase shifted animals shown by gene. (i) Phase difference (left) and absolute phase difference (right) are shown for memory and clock genes. (j) Cosinor fits for all genes are shown sorted based on time of peak expression. Red shows high expression, blue shows low expression. (k) Same as for controls but after forced desynchrony, sorted as in (j).

### Forced desynchrony abolishes circadian expression in many genes that affect memory

To assess rhythmicity, a COSINOR model with a fixed 24-hour period was fitted to each gene across time points, separately for each experimental group (resulting in 58 fits in total). To account for multiple hypothesis testing, p-values were adjusted using the false discovery rate (FDR) method^51^. A large proportion of the 29 genes had circadian gene expression in control animals, approximately 66 % (see Figures 2b, c, d, e, & k; supplementary Figures 2, 3, 4). In the shifted animals, the number of circadian genes was significantly reduced (p=0.03), with only 34 % of genes having circadian gene expression (Figure 2c). While many stopped having circadian gene expression, some genes maintained circadian rhythmicity, and a few genes developed circadian rhythmicity under forced desynchrony conditions (see Figure 2b).

### Amplitude was attenuated in almost all genes that displayed circadian rhythms

Amplitude of a circadian rhythm represents the maximum expression of the process being measured and when it’s changed there can be an impact on the function of that process. The majority of genes classified as circadian or suggestively circadian in controls displayed reduced amplitude in shifted animals, in many cases by more than 50% (Figure 2g; clock genes: *p* = 0.02; memory genes *p* = 0.002). The absolute changes in amplitude per gene subset were also significantly different from chance (Figure 2g & h; clock genes: *p* = 0.003; memory genes *p* < 0.001). The strongest reductions were observed for *Timeless*, *Rorα,* and *Per1*, with amplitude decreases of approximately 60%, 70%, and 80%, respectively. In contrast, *Bmal1* exhibited increased amplitudes under disruption, suggesting that some components of the circadian system may become more pronounced under altered environmental conditions.

### Phase changed in genes that remained circadian in disrupted animals

Phase of an oscillation refers to the time that maximal expression occurs (acrophase). This is an important aspect of a circadian oscillation because changes in phase mean that maximal and minimal times of expression are no longer aligned with the individual’s environment and uncoordinated changes across genes means that gene expression is no longer synchronized. This misalignment can thereby affect the function of processes dependent on the circadian rhythm. As demonstrated in Figures 2i & j, circadian disruption led to changes in phase, with the clock genes having a significant phase advance (*p* = 0.028). However, memory genes did not have a significant coordinated advance or delay (*p* = 0.388). Instead, memory genes showed uncoordinated changes in their expression, with both advances and delays in phase, shifting away from their usual peak expression time. Accordingly, absolute phase change was significant (memory genes, *p* = 0.002; clock genes, *p* = 0.028).

## Discussion

Circadian rhythm disruption in the form of six days of forced desynchrony changed the coordinated circadian expression of *clock* and memory genes. In controls, 66 percent of 29 genes from the Long Evans rat had probable circadian rhythms in expression. Most of the *clock* related genes – *Bmal1*, *Cry1*, *Cry2*, *Rorα, Per1, Per2, Dbt, Dbp, Timeless* displayed circadian or probable circadian rhythms in expression. Memory genes *Reln, Scop, Pka, Atf6b, Nrn1, Hdac3, CamkII, Mapk1, Dnmt3a, Sirt1* were also rhythmic. The clock gene findings are similar to mouse ^48,52^ and rat studies ^47,53,54^. This set of *memory* genes have not yet been assessed in one report this way, but studies in mice focusing on some of these genes in isolation have found rhythmicity ^11,13,14,30^.

After environmental circadian disruption, without up or down regulating absolute gene expression only 34 percent of 29 genes were rhythmic – *Reln, Scop, Pka, Dbp, Cry2, Per2, Bmal1, Clock, Dnmt3a,* and *Esr1.* Genes tended to have reduced amplitude of oscillation and those that remained rhythmic exhibited phase changes, with the *clock* genes displaying a significant phase advance and *memory* genes having a significant absolute phase change but not a coordinated one. Thus, after circadian rhythm disruption, molecular timing goes awry without up or down regulating absolute gene expression.

While we investigated acute disruption, chronic disruption has also been studied. Similar to our finding of several genes becoming circadian, in the liver many genes in the genome became rhythmic after exposure to a high fat diet, which also results in circadian rhythm disruption^55^. A paradigm that involved two light inversions every six days for 90 days in rats found memory impairments and changes in some of the core clock genes and genes involved in plasticity like glutamate, AMPA, and NMDA receptors in the hippocampus^47^. They, like others, analyzed differences at every ZT extraction point instead of globally determining if that gene is still rhythmic and if so, whether the amplitude or phase significantly change. These circadian characteristics were analyzed in a recent study that used a photoperiod shifting paradigm in mice, which found that most of the circadian genes-maintained expression patterns, whereas two of the seven plasticity genes (none assessed in the current study) rhythms were abolished^48^. The fact that the findings from the current study are arguably more deleterious than these 90-day paradigms stresses the efficacy of forced desynchrony and acute paradigms to disrupt the hippocampal circadian clock. In future work, the effects of acute and chronic circadian rhythm disruption paradigms should be addressed in the same investigation.

In conclusion, we report that when circadian rhythms are decoupled from their zeitgebers, there is not up or down regulation in hippocampal gene expression of core clock or memory genes but instead changes in the circadian expression of these genes. Notably many circadian genes stop oscillating when exposed to disruption. The findings suggest that changes in the oscillatory patterns of genes could be involved in the relationship between circadian rhythms and memory, which may be helpful for understanding disease states in which cognition and circadian rhythms are both disrupted (See Figure 3).

**Figure 3.**
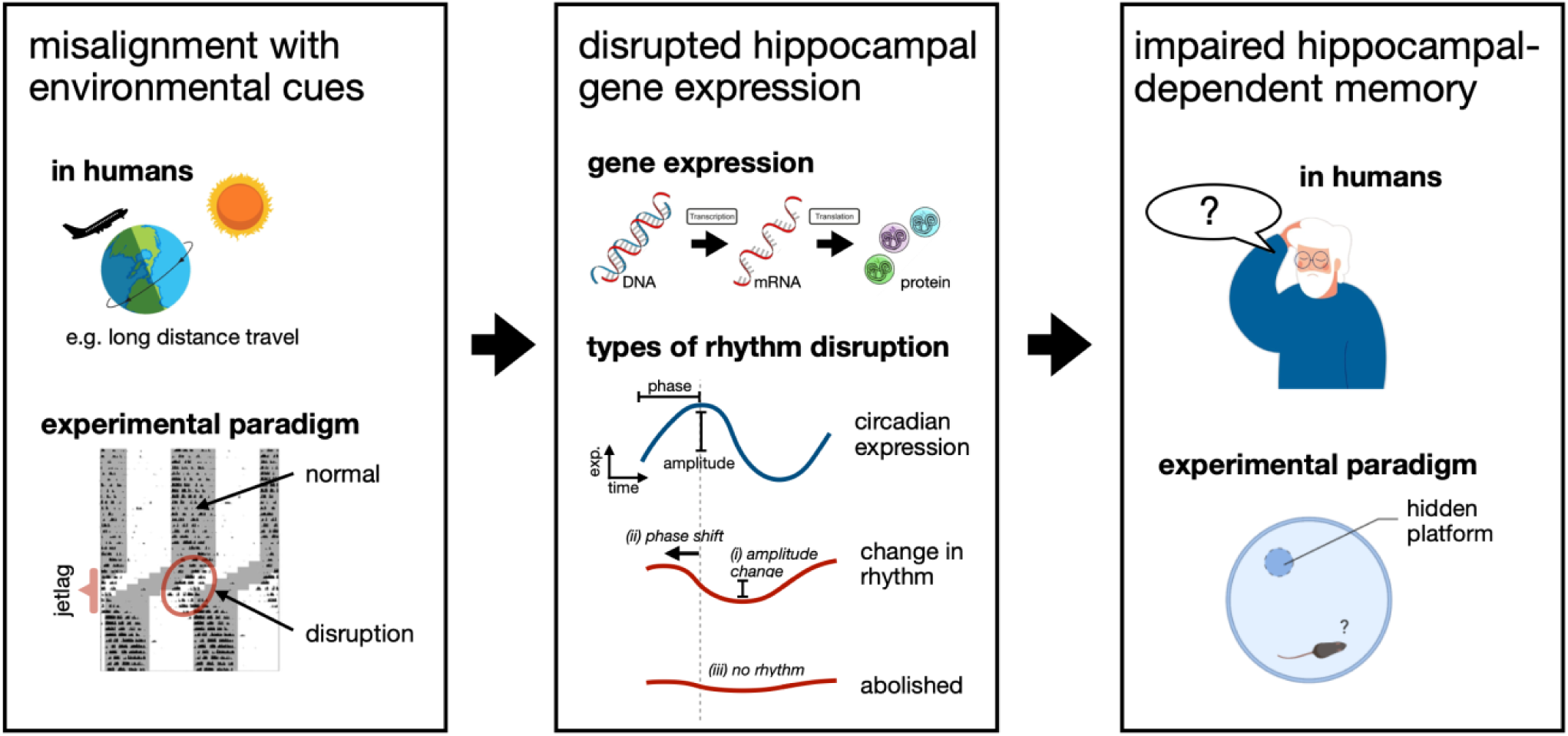
Circadian rhythm disruption likely affects memory via altered hippocampal oscillations in the expression of genes involved in the clock and memory. Panel 1) In humans, jetlag, shift work, night work, social jetlag, and disease states involve a misalignment of activity and sleep with that dictated by their biological clock. Our experimental paradigm mimics this, as during the six-days of three-hour phase advances (21-hour day) activity (shown in the actogram; shaded regions represent lights off) and sleep occur when they should not. There is much more activity when the animals should be sleeping, and much more sleep when the animals should be active. Panel 2) The current data and others find that genes involved in the generation of circadian rhythms and those involved in memory oscillate in the hippocampus. Genetic manipulation to some of the genes tested here like *Bmal1, Per1, and Sirt1* in whole brain or in just hippocampus impairs memory and the circadian properties of other clock and memory genes in the hippocampus. Our conclusions here that environmental circadian rhythm disruption paradigm alters hippocampal circadian rhythmic gene expression three different ways in a gene dependent manner are strengthened and extend the findings from genetic mouse models. i) the amplitude of the circadian rhythm changed for both clock and memory genes. ii) The time of maximal expression (phase) of circadian expression changed for genes involved in both the clock and memory. iii) For some clock and memory genes, they no longer met the criteria for a circadian rhythm following the 21-hour day paradigm. Panel 3) Circadian rhythm disruption associated with jetlag, altered work schedules, genetics, or various disease states, such as mental illnesses, natural aging, dementia, and Alzheimer’s disease, is also accompanied by impaired memory. Our 21-hour day rodent model of circadian rhythm disruption impairs long-term hippocampal dependent memory in the most typical task to assess its function (place version of the Morris water task). When integrated with the body of the literature in both humans and animal models, these findings may be relevant to the memory impairments seen in aging, dementia and Alzheimer disease.

## Methods

### Experimental Model and Subjects

48 male Long Evans rats approximately 90 days old from Charles River colonies (QC, Canada) were divided randomly into two groups of 24 animals (control: n = 24; shifted: n = 24). The animals for each group were further divided across six time points in a 24-hour period (each time point: control: n = 4; shifted: n = 4). Animals were pair-housed in clear Plexiglas hanging cages with beta-cob bedding, and a black tube. The colony rooms were kept at a temperature of 21◦ C and a humidity of 35%. Before implementation of the circadian rhythm disruption paradigm animals were allowed to acclimate to the colony room for two-weeks with a light dark cycle of 12L:12D. A red-light bulb (< 5 lux) remained on all the time. The rats always had *ad libitum* access to food and water. All experimental procedures were approved by the Canadian Centre for Behavioral Neuroscience, University of Lethbridge care committee and performed in accordance with the standards set out by the Canadian Council for Animal Care.

### Procedures

On Day one of the circadian rhythm disruption paradigm, the control group remained on a 12L:12D light dark cycle for six consecutive days, with lights turning off at 19:00 (see Figure 1). Rats in the circadian rhythm disruption group were exposed to a 21-hour day 9L:12D for six consecutive days, which is the equivalent to a three-hour phase advance each day for six consecutive days (see Figure 1). Home cage wheel access, nor behavioral memory training were used in the current study as they can affect circadian rhythms in activity and gene expression ^56,57^, thus leaving it impossible to determine the effects of the paradigm on hippocampal circadian gene expression. Nonetheless, this exact paradigm affects circadian rhythms, sleep, and hippocampal dependent memory ^20,21,24,58^.

On day seven, all rats were sacrificed every 4-hours across six timepoints. Four circadian rhythm disrupted rats, and four rats exposed to a 12L:12D lighting conditions were sacrificed at each time point. Zeitgeber time (ZT) refers to the timing of a stimulus, with ZT0 being the time that the colony lights came on (see Figure 1). Although phase changes in the shifted animals (circadian time differs from ZT time), period of activity and sleep are not affected by this forced desynchrony paradigm^20,24,58^. The rats were anesthetized with isoflurane, decapitated so the whole hippocampus could be extracted and then flash frozen with liquid nitrogen at ZT2, ZT6, ZT10, ZT14, ZT18, and ZT22, respectively. Illumina NextSeq500, single-end, multiplexed RNA sequencing was performed with 75 cycles. Illumina CASAVA 1.9 with default settings was used for base-calling and demultiplexing. Because base qualities were excellent across all samples and no adapter contamination was found, no adapter removal was performed. Subsequently, RNA sequencing was performed to quantify mRNA expression levels across the hippocampal transcriptome. The resulting raw count data for 14,789 genes were processed in R^59^, where data normalization and a variance stabilizing transformation (VST) were applied using the DESeq2 package^60,61^. During quality control, two rats from the circadian disrupted group were identified as outliers and excluded from all further analyses (one ZT18; one ZT22). This resulted in a final sample size of N = 46 (n = 24 in the control group; n = 22 in the shifted). Following outlier removal, the remaining data were renormalized and mapped using the Rattus norvegicus reference genome from the Ensembl database (http://useast.ensembl.org/Rattus_norvegicus/Info/Index).

### Quantification and Statistical Analysis Average expression

R ^59^ and Python ^62^ were used for all statistical analyses and figures. Average gene expression analysis was performed using the DESeq2 package in R^60^. This framework models gene expression count data using generalized linear models (GLMs) with a negative binomial distribution, thereby accounting for the overdispersion typically observed in RNA-seq datasets^60^. One possible modeling strategy is to treat each combination of time and condition as a single factor (e.g., ZT2 C) in the model. However, such a design does not explicitly separate time-of-day effects and treats each time point independently. As a result, it prevents the model from borrowing information across time points and reduces statistical power.

To address this limitation, we employed an additive design (∼ Time + Condition), which models the effects of circadian time and experimental condition separately. This approach first accounts for structured circadian variation in gene expression, allowing the model to isolate the effect of condition on top of this temporal baseline. In this framework, comparisons between conditions are effectively made at matched time points, while all samples collectively contribute to estimating a time-averaged condition effect. Following model fitting, we obtained estimates of log2 fold changes (LFCs) along with corresponding raw p-values and false discovery rate (FDR)-adjusted q-values for each gene. To improve the reliability of these estimates, particularly for genes with low counts or high variability, we applied shrinkage of log2 fold changes using the lfcShrink function^63^. This method moderates extreme fold-change estimates by borrowing information across genes, without altering statistical significance. The resulting differential expression profiles were visualized using a volcano plot of the shrunken log2 fold changes against the -log10(FDR). Genes exhibiting both large effect sizes and strong statistical significance appear in the upper corners of the plot (see Figure 1).

### Cosine Fit

To assess rhythmicity, a COSINOR model with a fixed 24-hour period was fitted to each gene across time points, separately for each experimental group (resulting in 58 fits in total). Prior to model fitting, gene expression values were Log-transformed and standardized (z-scored) by subtracting the overall mean (across both groups) and dividing by the corresponding standard deviation.

The COSINOR model included two key parameters: amplitude and phase. Model parameters were estimated using weighted least squares, allowing us to account for heteroskedasticity observed across time points. Model fit was evaluated using a weighted regression F-test, with a significance threshold of *α* = 0.05.

To account for multiple hypothesis testing, p-values were adjusted using the false discovery rate (FDR) method^51^. In addition, the corrected Akaike Information Criterion (AICc) was computed and then converted into P(rhythmic), an Akaike weight quantifying the relative support for the rhythmic COSINOR model compared with a constant-expression null model. Values closer to 1 indicate stronger support for rhythmic (wave) expression, whereas values closer to 0 indicate support for the null model.

Genes were classified as circadian if they satisfied both criteria of *q* − *value <* 0.05 and *P*(*rhythmic*) *>* 0.8. Genes were labeled as suggestively circadian if they exhibited *P*(*rhythmic*) *>* 0.5 and q-value < 0.1. All remaining genes were classified as non-circadian. A Fisher’s exact test was used to compare if the fraction of genes that were classified as circadian or suggestively circadian in the control animals changed in the circadian disrupted animals.

Of note, is that with the COSINOR model we are asking how well normal 24-hour rhythms explain the data. For shifted animals, we are asking whether their normal 24-hour rhythms which are coordinated across genes remain coordinated despite the unentrainable LD cycle.

### Amplitude Analysis

For the amplitude analysis, we included genes classified as circadian or suggestively circadian in at least one of the two conditions. Amplitude estimates were recalculated using the non-standardized, log-transformed expression data to preserve the original scaling of the measurements. Subsequently, both raw and absolute differences in amplitude between conditions were computed. To evaluate significant changes in raw and absolute amplitudes, we performed one-sample t-tests across three distinct gene groupings: all genes combined, clock genes and memory genes.

### Phase Analysis

Genes that met at least one of the two criteria for circadian or suggestively circadian rhythmicity in both conditions were further analyzed to assess potential phase shifts. Phase estimates were obtained from the COSINOR model fits for each gene in each group, and these were used to calculate both raw and absolute phase differences between conditions. Differences were wrapped to (−12, +12) hours to account for 24-hour circularity, ensuring the shortest phase distance was used. To evaluate potential raw and absolute phase differences, we performed a one-sample t-test against zero across the same three distinct groupings: all genes combined, clock genes and memory genes.

## Resource Availability

Main contacts: Drs. Robert McDonald and Scott Deibel

Data availability: A dataset with the expression values of the genes reported above from unilateral hippocampal tissue in the Long-Evans rats exposed to a 12L:12D light dark cycle (n = 24) and a 21-hour day (n = 22; 9L:12D) is being freely provided. Hippocampus was extracted on day seven, every four hours, after six days of exposure to either a 24-hour day or a 21-hour day. This paradigm disrupts circadian rhythms, sleep, and impairs memory. ZT refers to zeitgeber time, with ZT0 representing when the colony lights turn on.

The dataset and complete code for analyses are under embargo at figshare and can be accessed via the link below.

https://figshare.com/s/3a03395842661be6db25

## Acknowledgments

We thank Yaroslav Ilnytskyy, Andrey Golubov, and David Z. Kochan for their help in tissue processing and analysis. We thank the members of the McDonald, and Kovalchuk laboratories for their guidance, and support for this project.

## Author Contributions

Conceptualization: SHD, RJM, ABL

Methodology: SHD, RJM, OK, ABL, CB, SM, EF

Investigation: SHD, NSH, OK Analysis: ABL, SM, SHD

Visualization: ABL, SHD, SM, EF, IH

Supervision: RJM, SHD, ABL

Writing—original draft: SHD

Writing—review & editing: all authors

## AI Statement

AI was not used in any aspect of the planning, experiment, analysis, or writing process.

---

**Supplementary Figure 1.**
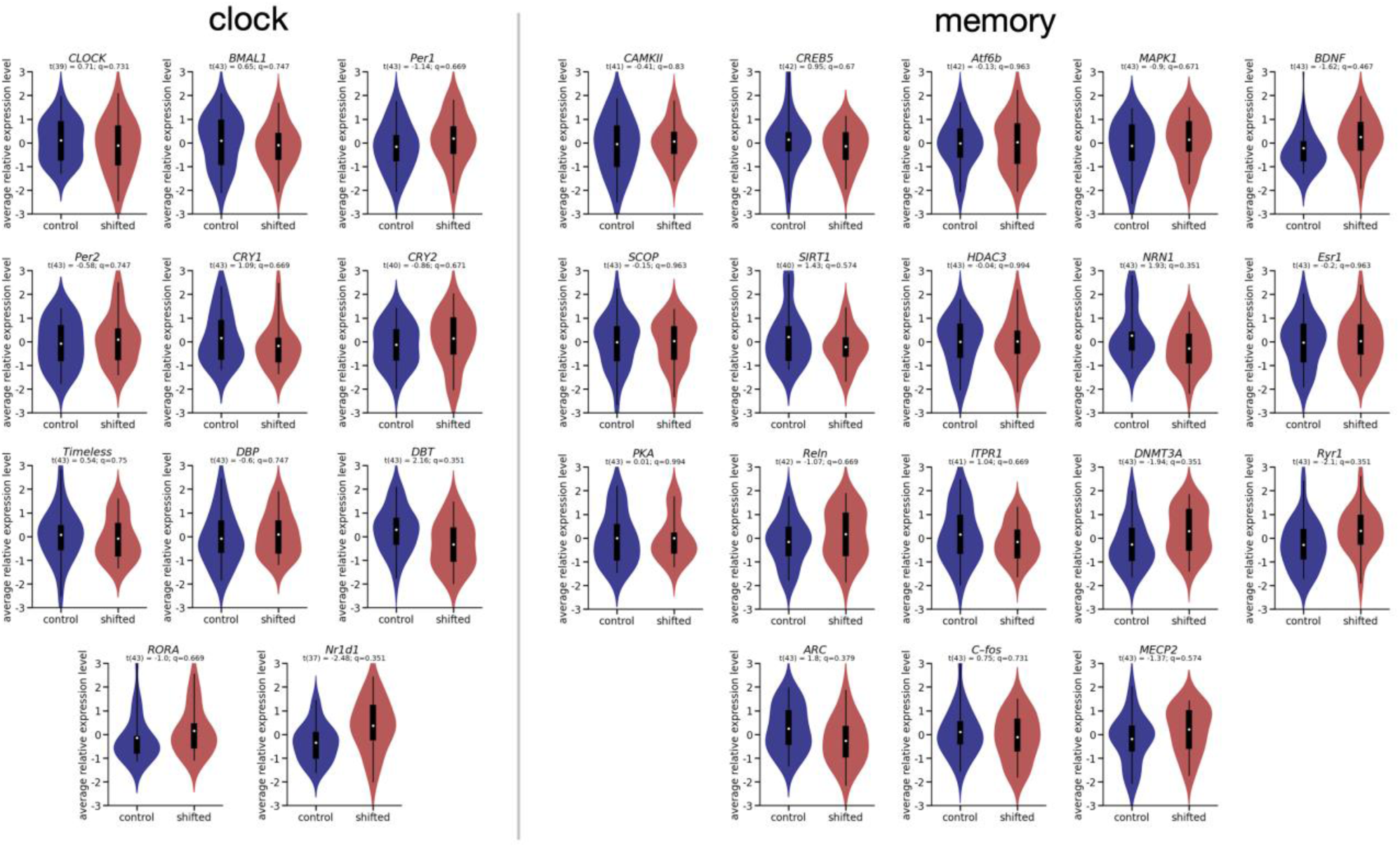
Average expression of *clock* and memory genes for control and circadian rhythm disrupted animals.

**Supplementary Figure 2.**
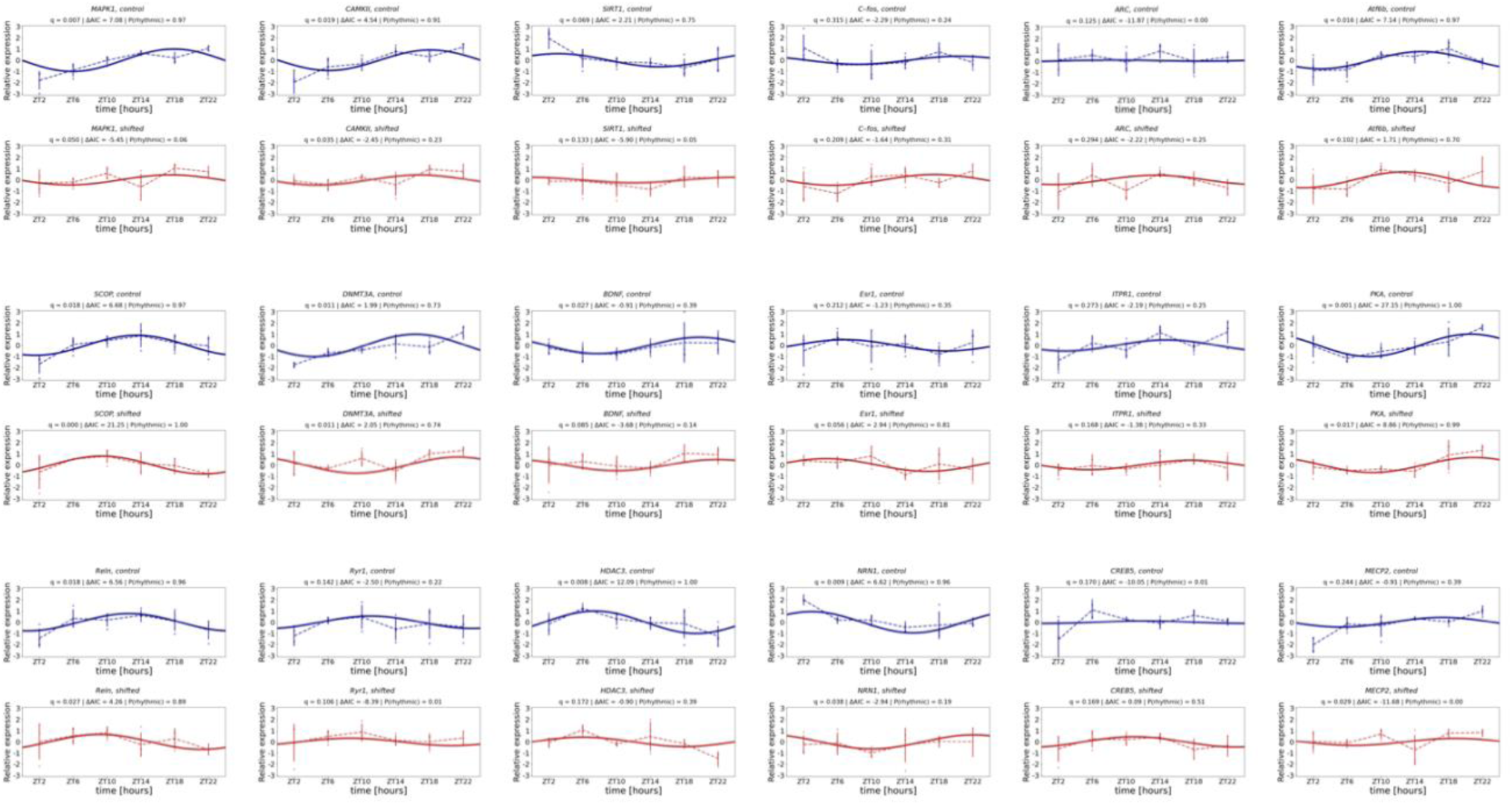
Cosine fits of memory genes in control and circadian rhythm disrupted animals.

**Supplementary Figure 3.**
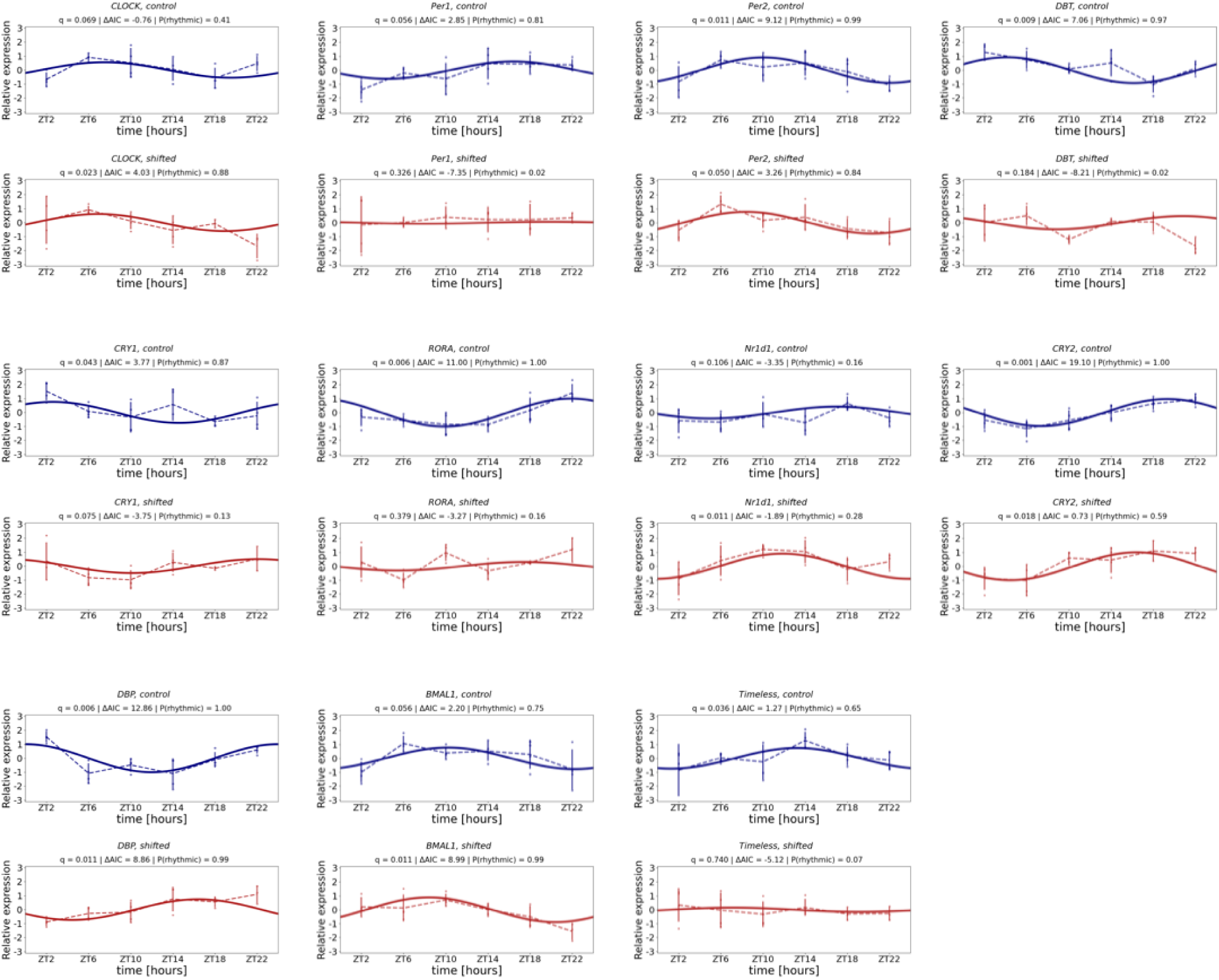
Cosine fits of *clock* genes in control and circadian rhythm disrupted animals.

**Supplementary Figure 4.**
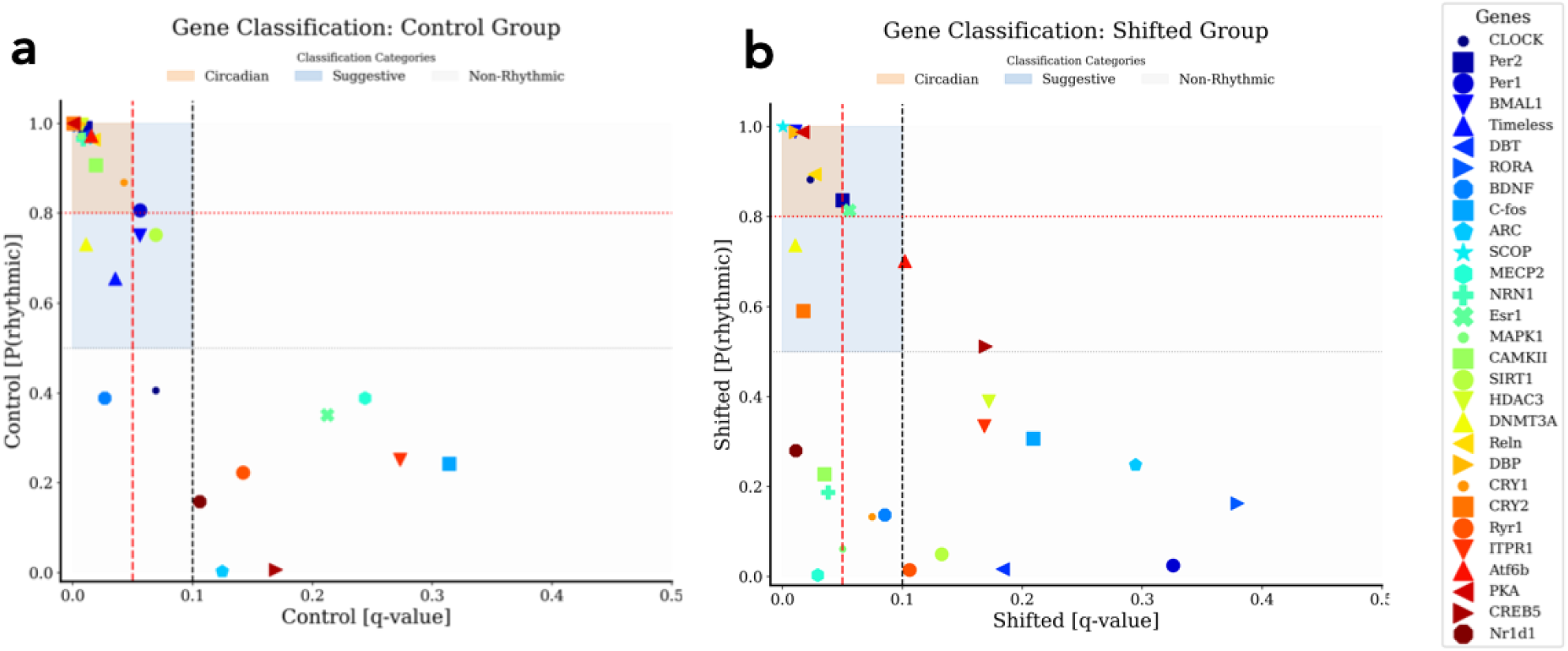
Criteria for selecting circadian genes. See methods for explanation of criteria.

